# Precautionary Coherence Unravels Dose Escalation Designs

**DOI:** 10.1101/240846

**Authors:** David C. Norris

## Abstract

**Background:** Coherence notions have a long history in statistics, as rhetorical devices that support the critical examination of statistical doctrines and practices. Within the special domain of dose-finding methodology, a widely-discussed coherence criterion has been advanced as a means to guard the conceptual integrity of formal dose-finding designs from *ad hoc* tinkering. This is not, however, the only possible coherence criterion relevant to dose finding. Indeed, a new coherence criterion emerges naturally when the near-universal practice of cohort-wise dose escalation is examined from a clinical perspective.

**Methods:** The practice of enrolling drug-naive patients into an escalation cohort is considered from a *realistic* perspective that acknowledges patients’ heterogeneity with respect to pharmacokinetics and pharmacodynamics. A new coherence criterion thereby emerges, requiring that an escalation dose be tried preferentially in participants who have already tolerated a lower dose, rather than in new enrollees who are drug-naive. The logical implications of this ‘precautionary coherence’ (PC) criterion are worked out in the setting of a 3+3 design. A ‘3+3/PC’ design that satisfies this criterion is described and visualized. A simulation study is performed, evaluating the long-run performance of this new design, relative to optimal 1-size-fits-all dosing.

**Results:** Under the PC criterion, the 3+3 dose-escalation design necessarily transmutes into a dose *titration* design. Two simple rules suffice to enable abandonment of low starting doses, and termination of escalation. The process of conducting the 3+3/PC trial itself models the application of a *dose titration algorithm* (DTA) that carries over readily into clinical care. The 3+3/PC trial also yields an interval-censored ‘dose-survival curve’ having a semantics that should prove familiar to oncology trialists. Simulated 3+3/PC trials yield DTAs over a median of 6 dose levels, achieving 50% improved population-level efficacy compared to optimal 1-size-fits-all dosing.

**Conclusions:** Dose individualization can be accomplished within a trial conducted along ‘algorithmic’ lines resembling those of the inveterate 3+3 design. The dose-survival curve arising from this ‘3+3/PC’ design has semantics that should prove familiar and conceptually accessible to oncology trialists, and also seems capable of supporting more formal statistical treatments of the design. In the presence of sufficient heterogeneity in individualized optimal dosing, a 3+3/PC trial outperforms *any conceivable* 1-size-fits-all dose-finding design. This fact eliminates the rationale for the latter designs, and should put an end to the further development and promulgation of 1-size-fits-all dose finding.

## INTRODUCTION

Axioms of coherence have long played a role in the polemics of the Bayesian vs Frequentist debate. As (Robins and Wasserman 2000) note, “Usually, these arguments lead to conclusions of the form that inferences are coherent if and only if they are Bayesian" Within the field of dose-finding methodology, (Cheung 2005) has introduced a notion of coherence that addresses a different tension—one that pits biostatisticians who strive for conceptually intact trial designs against clinicians who may feel compelled to make *ad hoc* modifications to dose-finding trials on the fly. Although commonly discussed without qualification simply as “coherence" (Wheeler 2016; Iasonos et al. 2016; Bartroff and Leung Lai 2011), the particular notion introduced by (Cheung 2005) is by no means the only possible such criterion in dose-escalation trials, nor even the most pertinent.

## PRECAUTIONARY COHERENCE

In any of the standard dose-escalation designs, when a dose escalation occurs it is common practice to enroll new, *drug-naive* participants at the escalated dose. Remarkably, this holds true even when the dosing interval has elapsed for some earlier-enrolled participants who *have already tolerated a lower dose of the drug* and are ready to receive their next dose. If one of these drug-naive participants were to experience a highly morbid or even fatal dose-limiting toxicity (DLT) at this first-in-human escalation dose, this practice might well appear incoherent in retrospect. Under any realistic conception of the heterogeneity of toxicity, participants who have already tolerated lower doses of the study drug must be presumed less likely than a drug-naive participant to experience a severe DLT.

These considerations lead naturally to a principle I term ‘precautionary coherence’:

> Escalation doses should be tried preferentially in participants who have already tolerated lower doses of the study drug, as opposed to newly-enrolled, drug-naive participants.

## THE 3+3/PC DESIGN

To obtain the most straightforward working-out of this principle, I will treat the case where the DLT assessment period is some integer multiple of the dosing period. (Without loss of generality, this can be regarded as equivalent to the specific case where DLT assessment and dosing both occur in discrete time with period 1, the latter process being *adapted to* the former.) The result of this simplifying assumption is that, at the time of any dose escalation decision *all enrolled participants are available to test the new dose*.

One appreciates immediately that precautionary coherence (PC) entails what is often called ‘intra-patient dose escalation’ (Simon et al. 1997; Dancey, Freidlin, and Rubinstein 2006), but which I will here call simply *dose titration*.^1^ Under this simplified terminology, a dose-escalation trial, when modified to achieve precautionary coherence, necessarily becomes a *dose titration* trial. More plainly: *a PC trial conducts escalation only through titration*.

Unlike a dose-escalation trial, which carries forward a *single* dose that serves as a current working hypothesis about ‘the’ MTD, a PC trial carries forward *two* doses: (a) a low ‘enrolling’ dose at which newly-enrolled participants begin their titration, and (b) a maximum dose beyond which titration is not pursued. The design of a dose titration trial must specify how each of these titration limits evolves as the trial proceeds and information accrues.

To see how PC might work out concretely in the context of a 3+3 design, we posit the following dose-dropping and escalation-stopping rules:

- **PC rule:** A yet-untried dose level *D* + 1 is administered only to participants who have *already* tolerated dose level D. This is logically equivalent to the dictum, “escalation only through titration”
- **cohort rule:** Escalation (*i.e.*, upward titration to a yet-untried dose) is performed only once a cohort of 3 or more participants has accumulated, who satisfy the PC rule.
- **reduction rule:** A participant who experiences a DLT at dose level *D* > 1 is thereafter continued at level *D* − 1. A participant who experiences a DLT at the lowest dose level *D* = 1 discontinues treatment.
- **exit convention:** Once a participant’s steady dose is determined, the participant is considered to have ‘exited’ the titration study, notwithstanding that (per the *reduction rule*) the participant continues to receive treatment at that steady dose.
- **bypass rule:** When, with 90% confidence, the current ‘enrolling’ dose proves tolerable to over 80% of participants, the enrolling dose is bumped up to the next dose level. (The bypassed low dose is however retained for dose reductions as needed.)
- **stop rule:** When, with 90% confidence, the highest dose proves tolerable by under 1/3 of participants, escalation stops. This means that no higher doses will be considered for the remainder of the study.
- **rollback rule:** When, with 90% confidence, the highest dose proves tolerable by under 1/4 of participants, this highest dose is abandoned for further titration. Any participants who have already tolerated this dose, however, are maintained on it without a dose reduction.

Before proceeding to examine *necessity* and other logical relations between these rules, we develop a more concrete understanding of them by visualizing a simulated trial.

## A SIMULATED TRIAL

To simulate a 3+3/PC trial, we suppose that MTD*_i_* is Gamma-distributed with coefficient of variation 0.7 as in (Norris 2017b) and with mean 1. Fixed dose levels are established in a geometric series starting at 0.25 and increasing by 40% at each step. A simulated trial is shown in Figure 1.

**Figure 1.**
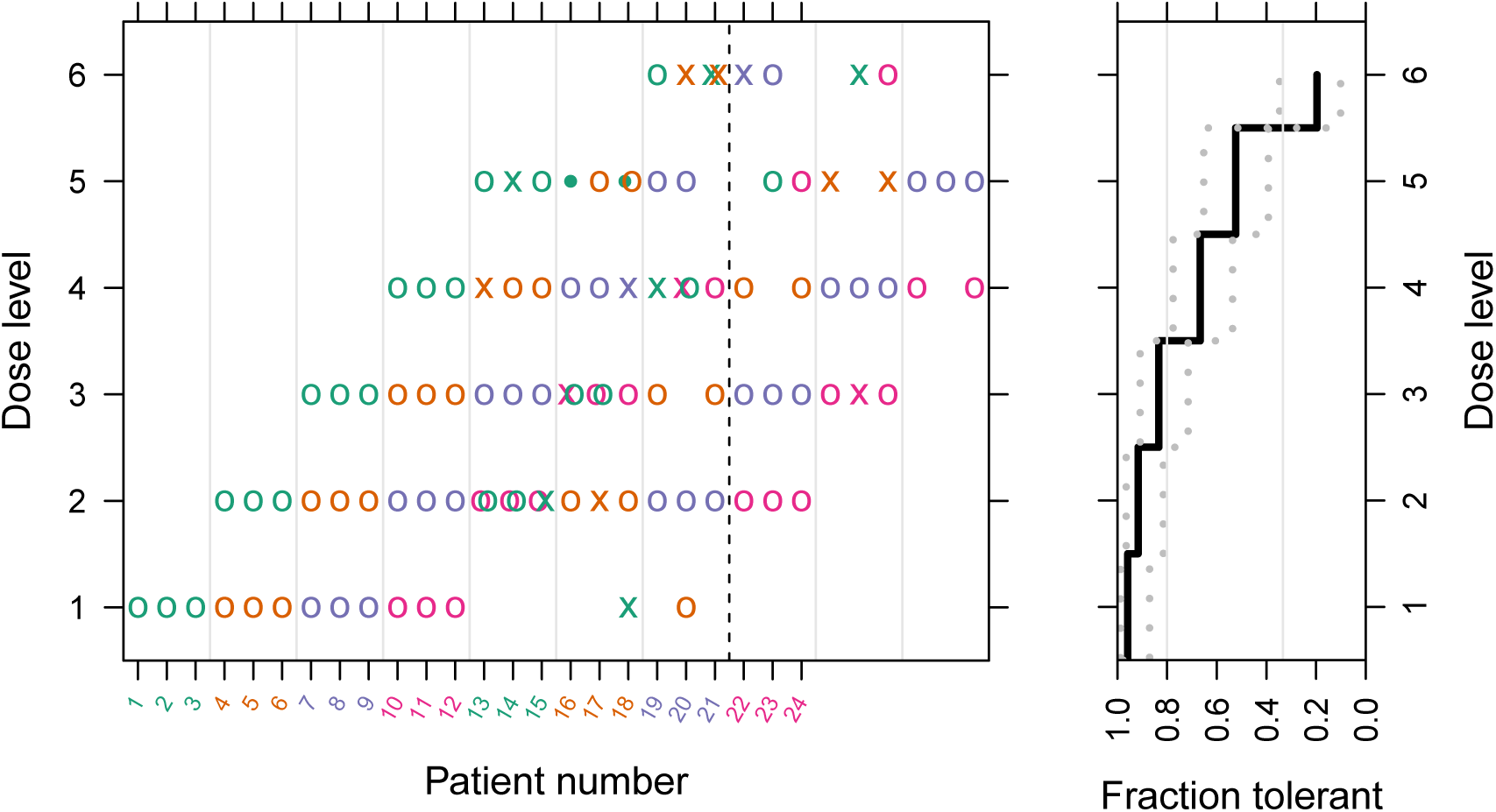
Simulated 3+3/PC study. Numbered participants 1–24 enroll 3-at-a-time into sequenced *DLT assessment periods*, and percolate through the ‘OX plot’ on the left until they exit titration with an interval-censored MTDj determination. The ‘dose-survival plot’ on the right depicts the accumulated interval-censored dose tolerance using the familiar semantics of the Kaplan-Meier curve, but with *dose* substituted for *time*. Left-right positioning within the DLT assessment periods, plus a cycle of 4 (colorblind-friendly) colors, facilitate tracing of individuals from enrollment to exit. For example, participants 1–3 enroll in period 1 at dose level 1, and titrate upward steadily to dose level 5 in period 5. Here, id2 experiences a DLT (denoted ‘x’) and exits the titration study with the determination MTD_2_ ∈ [4,5). The *cohort rule* not being satisfied, id1 and id3 hold at the same dose into period 6 (depicted ‘•’). With the dose escalation in the subsequent period 7, id3 exits with MTD_3_ ∈ [5,6). Because the *stop rule* triggers at the end of period 7 (indicated by the dashed vertical line), id1 also exits titration with an unbounded MTD1 G [6, ∞). After the *bypass rule* has triggered, as happens at the end of period 4 in this simulation run, participants may percolate *downward* through the OX plot. For example, id15 and id17 are seen to experience DLTs upon enrollment, and so undergo dose reductions; they exit at the end of periods 6 and 7, respectively, with determinations MTD_15_ ∈ [0,1) and MTD_17_ ∈ [1,2). An 80% confidence band is shown on the dose-survival curve, as constructed by R package km.ci (Strobl 2009) using a method due to (Rothman 1978). Thresholds corresponding to the *bypass* and *stop* rules are also marked with vertical lines. It may be instructive to observe that the dose-survival curve as shown would trigger bypass of dose level 2, and that it would (just barely) *not* trigger the stop rule. (These are moot points at the end of period 10, however, since enrollment ended with period 8, and the ‘stop’ question was already decided at the end of period 7.)

## ON THE NECESSITY OF THE 3+3/PC RULES

The 3+3/PC trial demonstrated here departs so dramatically from the familiar 3+3 design, that the rules set forth above might be supposed each to have made an independent contribution to this departure. Indeed, only the *cohort rule* appears to preserve anything of the spirit of the 3+3 trial. Nevertheless, my view is that these rules constitute a straightforward working-out of the logical implications of precautionary coherence. I believe that any PC dose-finding trial that preserves a ‘cohort’ concept like that embodied in the *cohort rule* (and is conducted over a predetermined discrete set of dose levels for a single drug) must necessarily adopt the remaining rules.

To see how this is so, suppose we adopt the PC and cohort rules, and examine the remaining rules in order. The *reduction rule* expresses nothing more than our ethical responsibility to trial participants which becomes immediately apparent in any dose titration design^2^. The *exit convention*, considered from the most mundane perspective^3^, becomes necessary for decluttering the OX plot on the left of Figure 1. Considered from a more refined perspective, however, an ‘exit’ concept does seem to arise naturally as a corollary to the *reduction rule* and its associated ‘terminal state’. In this state, the participant’s MTD*_i_* interval ceases to shrink, and the participant consequently ceases to contribute new information to the study's dose-survival curve. This curve itself seems indeed an inevitable construct, once one has decided to obtain information about a continuously distributed MTD*_i_* (Norris 2017a) through titration over a discrete set of doses.

Finally, once the dose-survival curve emerges, it constitutes the natural basis for making the *bypass*, *stop*, and *rollback* decisions. I make no strong claims as to the inevitability of the exact manner in which I have specified the latter 3 rules, however; these surely can be readily improved.

## PERFORMANCE CHARACTERISTICS OF 3+3/PC

We simulate 1000 3+3/PC trials like that of Figure 1. Each trial enrolls *N* = 24 participants with randomly-drawn MTD*_i_* ~ Gamma(*α* = 0.7^−2^, *β* = *α*)^4^, uses a discrete set of dose levels in a geometric sequence 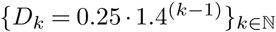, and runs for a total of 10 DLT assessment periods. The final (period-10) dose-survival curve from each simulated study is taken to define the recommended *initial dose* and *maximum dose* as per the bypass and stop rules, respectively. Results are shown in Figure 2.

**Figure 2.**
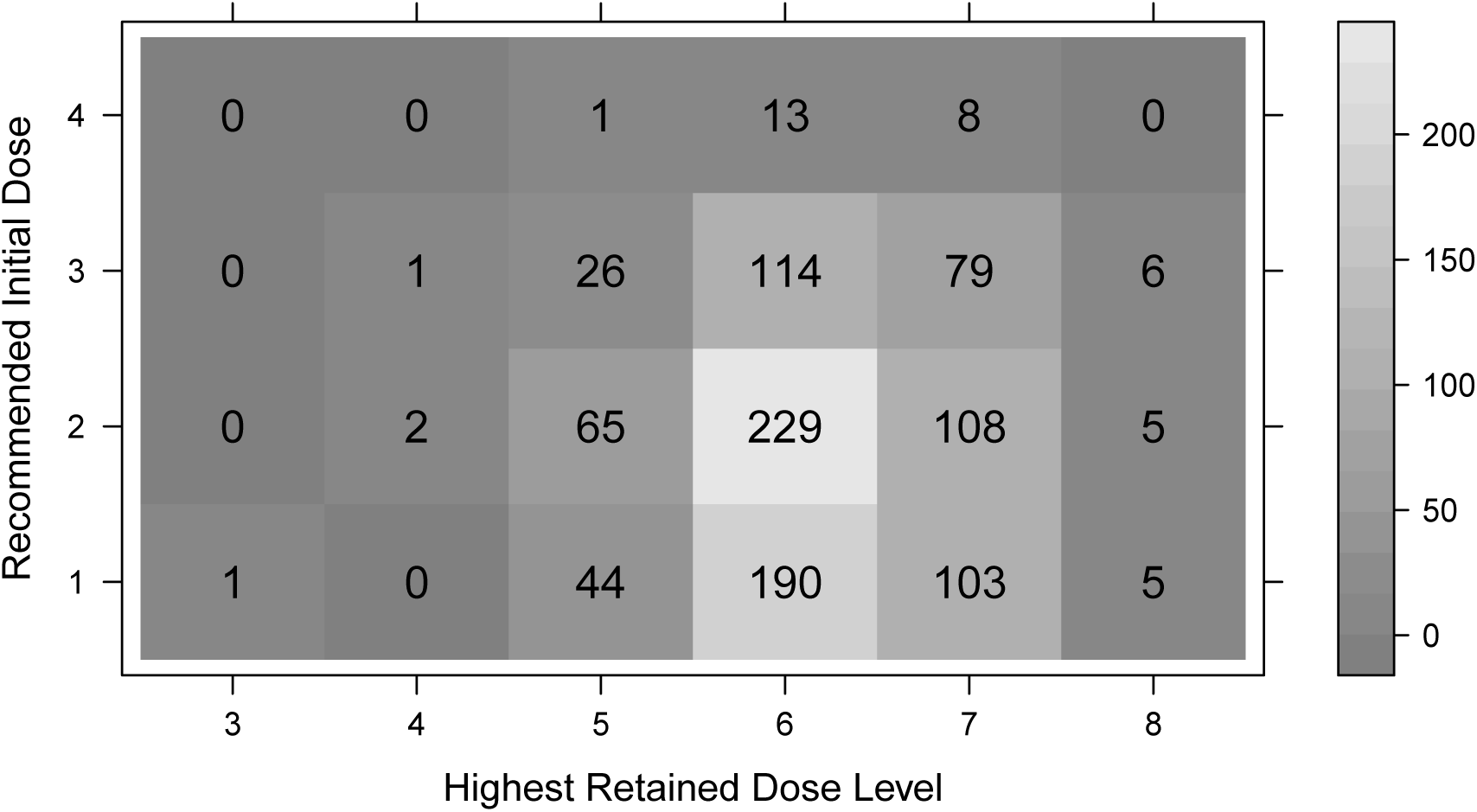
Joint frequencies of recommended *initial* and *maximum* doses from 1000 simulated 3+3/PC studies. Each simulated trial enrolls 24 participants with randomly drawn MTD*_i_* ~ Gamma(*α* = 0.7^−2^, *β* = *α*), employs a fixed set of dose levels in a geometric sequence *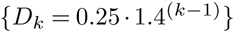*, and runs for 10 DLT assessment periods. Determinations of recommended initial and maximum doses are made from end-of-study dose-survival curves, according to the *bypass* and *stop* rules defined in the text. The modal trial recommends dose level 2 as starting dose for an individual-patient titration, and dose level 6 as a maximum. More than 95% of the simulated trials yield recommendations within ±1 of these modal levels.

Provided that we can safely titrate over the retained doses to choose the best one for each individual patient, then it makes sense to estimate a population-level efficacy according to (Norris 2017b, Equation 12). Table 1 shows the efficiency (relative to perfect, individualized MTD*_i_* dosing) for 1-size-fits-all dosing at each of dose levels 1–7, as well as for titration over increasing subsets of these levels up to the full set of 7. Of the doses considered, level 2 proves near-optimal for 1-size-fits-all dosing. Titration over the modal retained 6 dose levels (see Figure 2) achieves more than 80% efficiency, a better-than 50% improvement upon 1-size-fits-all dosing in this scenario.

**Table 1.** Efficiency of 3+3/PC dose titration vs 1-size-fits-all dosing. The 7 dose levels considered range from the 9th to 91st percentiles of MTD*_i_* in the simulated population. One-size-fits-all dosing achieves its peak efficiency (53%) near dose level 2. The majority of the efficiency gains from titration are realized using the first 5 dose levels. Effiencies are quoted relative to perfect ‘MTD*_i_* dosing’.

| Dose level | Dose (mg) | $MTD_i$ Quantile | 1-size-fits-all | Titration |
| --- | --- | --- | --- | --- |
| 1 | 0.2500 | 0.0888 | 0.5037 | 0.5037 |
| 2 | 0.3500 | 0.1549 | 0.5257 | 0.5851 |
| 3 | 0.4900 | 0.2577 | 0.5127 | 0.6645 |
| 4 | 0.6860 | 0.4026 | 0.4521 | 0.7345 |
| 5 | 0.9604 | 0.5812 | 0.3443 | 0.7878 |
| 6 | 1.3446 | 0.7635 | 0.2121 | 0.8207 |
| 7 | 1.8824 | 0.9071 | 0.0965 | 0.8356 |

## DOES TITRATION MAKE DLT’S INEVITABLE?

Interpreted literally, the titration scheme described here will cause DLTs in all patients except those whose MTDj’s exceed the top dose. This highlights the importance of modifying the titration procedure in line with a realistic, *graded* conception of toxicity (Norris 2017c). The ‘intrapatient escalation option B’ of (Simon et al. 1997) illustrates this idea, stopping upward titration once a moderate toxicity is observed.

## DISCUSSION

As elements in a *dose individualization* design, the triplet cohorts inherited by 3+3/PC do seem egregious. Certainly, model-based PC designs can and should dispense with these triplets, to enable enrollment and titration of participants *singly*—as befits a dose individualization study. I do hope such designs will be forthcoming from the community of dose-finding methodologists. If, as I hope, the further development of 1-size-fits-all dose-finding methods ceases altogether, the talents of a large number of methodologists may be fruitfully redirected to such work.

The 3+3/PC design presented here serves 2 functions in relation to this hoped-for revolution in dose-finding methodology. The first is the *critical* function of demonstrating that:

> 3+3 is *not* the problem; 1-size-fits-all dose finding is the problem!

This point is made most conspicuously by my demonstration that a 3+3-type titration design can outperform even *optimal* 1-size-fits-all dosing—and consequently *any conceivable* 1-size-fits-all dose-finding design.

The second function of 3+3/PC is to serve as the basis for fielding pragmatic dose-individualization designs pending the development of more sophisticated, model-based designs. Notwithstanding its significant departures (noted above) from the standard 3+3 dose-escalation design, 3+3/PC retains aspects of the ‘3+3 spirit’ that account for its longstanding popularity. Chief among these are its apparent freedom from complex modeling assumptions, and its ‘transparency’ (Ji and Wang 2013). Even the most sophisticated element of 3+3/PC—the dose-survival curve—follows an idiom that oncology trialists should find utterly familiar. Until credible, model-based alternatives emerge, the 3+3/PC design may itself sustain further development in various directions; the problem of *cumulative toxicities* points in one such direction.^5^

The *precautionary coherence* principle introduced here also serves a primarily *critical* function, which it achieves with 3 distinct audiences. The term ‘coherence’ speaks in no uncertain terms to the biostatisticians who are chiefly responsible for the current crop of 1-size-fits-all dose finding methodologies. This audience will appreciate that their designs are undeniably *incoherent* in this trenchant sense. The term ‘precautionary’, on the other hand, speaks to clinical trialists’ overriding concern for conducting safe and ethical trials. For this audience, PC casts the apposite deficiencies of dose-escalation designs in a harsh and unflattering light. Finally, to those responsible for oversight of the clinical trials enterprise—especially the increasingly sophisticated community of patient advocates—‘PC’ encapsulates an important clinical intuition so as to make it an effective critical tool even in the hands of laypersons.

## CONCLUSIONS

I have introduced ‘precautionary coherence’, a new coherence criterion applicable to dose-finding trials, and demonstrated that it entails (individualized) dose *titration* as against (groupwise) dose *escalation*. The logical implications of precautionary coherence are worked out in detail within the setting of a 3+3-style trial. A transparent visualization, incorporating a ‘dose-survival curve’ with familiar Kaplan-Meier semantics, aids in the exposition of the resulting ‘3+3/PC’ design. Simulation of this design demonstrates its markedly superior population-level efficacy compared to 1-size-fits-all dose finding. The simplicity and transparency of 3+3/PC may help *dose individualization* to gain a foothold in early-phase trials, pending the availability of more formal, model-based methodologic developments.

## DATA AVAILABILITY

*Open Science Framework:* The figures and tables in this paper may be reproduced using R package DTAT (v0.2–1) together with code file pc.R, both available at doi:10.17605/osf.io/8n3pf.

### Grant information

The author declared that no grants were involved in supporting this work.

## Footnotes

1 Compare language employed by (Senn 2007, 318) contrasting “between-patient group escalations” against “within-patient titrations.”

2 It is interesting to contemplate how the expediencies of groupwise dose-escalation designs serve to obscure these responsibilities.

3 I will confess, this is how it first occurred to me.

4 This gives the MTD*_i_* distribution a coefficient of variation (CV) of 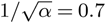, and a mean of *α/β* = 1.

5 The *variance reduction* achieved by titrating to individualized doses that approximate each participant’s MTD*_i_* may well serve to accelerate the detection of cumulative toxicities, and improve inference about them.

## REFERENCES

Bartroff, Jay, and Tze Leung Lai. 2011. “Incorporating Individual and Collective Ethics into Phase I Cancer Trial Designs.” Biometrics 67 (2): 596–603. doi:10.1111/j.1541-0420.2010.01471.x.

Cheung, Ying Kuen. 2005. “Coherence Principles in Dose-Finding Studies.” Biometrika 92 (4): 863–73. doi:10.1093/biomet/92.4.863.

Dancey, Janet, Boris Freidlin, and Larry Rubinstein. 2006. “Ch 4: Accelerated Titration Designs.” In Statistical Methods for Dose-Finding Experiments, edited by S. Chevret, 91–113. John Wiley & Sons, Ltd. http://onlinelibrary.wiley.com/doi/10.1002/0470861258.ch4/summary.

Iasonos, Alexia, Nolan A. Wages, Mark R. Conaway, Ken Cheung, Ying Yuan, and John O’Quigley. 2016. “Dimension of Model Parameter Space and Operating Characteristics in Adaptive Dose-Finding Studies.” Statistics in Medicine 35 (21): 3760–75. doi:10.1002/sim.6966.

Ji, Yuan, and Sue-Jane Wang. 2013. “Modified Toxicity Probability Interval Design: A Safer and More Reliable Method Than the 3 + 3 Design for Practical Phase I Trials.” Journal of Clinical Oncology: Official Journal of the American Society of Clinical Oncology 31 (14): 1785–91. doi:10.1200/JC0.2012.45.7903.

Norris, David C. 2017a. “Dose Titration Algorithm Tuning (DTAT) Should Supersede ‘the’ Maximum Tolerated Dose (MTD) in Oncology Dose-Finding Trials.” F1000Research 6 (July): 112. doi:10.12688/f1000research.10624.3.

Norris, David C. 2017b. “Costing ‘the’ MTD.” bioRxiv, August, 150821. doi:10.1101/150821.

Norris, David C. 2017c. “One-Size-Fits-All Dosing in Oncology Wastes Money, Innovation and Lives.” Drug Discovery Today, November. doi:10.1016/j.drudis.2017.11.008.

Robins, James, and Larry Wasserman. 2000. “Conditioning, Likelihood, and Coherence: A Review of Some Foundational Concepts.” Journal of the American Statistical Association 95 (452): 1340–6. doi:10.1080/01621459.2000.10474344.

Rothman, K. J. 1978. “Estimation of Confidence Limits for the Cumulative Probability of Survival in Life Table Analysis.” Journal of Chronic Diseases 31 (8): 557–60.

Senn, Stephen. 2007. Statistical Issues in Drug Development. 2nd ed. Statistics in Practice. Chichester, England; Hoboken, NJ: John Wiley & Sons.

Simon, R., B. Freidlin, L. Rubinstein, S. G. Arbuck, J. Collins, and M. C. Christian. 1997. “Accelerated Titration Designs for Phase I Clinical Trials in Oncology.” Journal of the National Cancer Institute 89 (15): 1138–47.

Strobl, Ralf. 2009. Km.ci: Confidence Intervals for the Kaplan-Meier Estimator. https://CRAN.R-project.org/package=km.ci.

Wheeler, Graham M. 2016. “Incoherent Dose-Escalation in Phase I Trials Using the Escalation with Overdose Control Approach.” Statistical Papers, June, 1–11. doi:10.1007/s00362-016-0790-7.

